# Microtubule tyrosination/detyrosination specifies a mitotic error code

**DOI:** 10.1101/801977

**Authors:** Luísa T. Ferreira, Bernardo Orr, Girish Rajendraprasad, António J. Pereira, Carolina Lemos, Joana T. Lima, Clàudia Guasch Boldú, Jorge G. Ferreira, Marin Barisic, Helder Maiato

**Affiliations:** Chromosome Instability & Dynamics Group, i3S - Instituto de Investigação e Inovação em Saúde, Universidade do Porto, Rua Alfredo Allen 208, 4200-135 Porto, Portugal; Instituto de Biologia Molecular e Celular, Universidade do Porto, Rua Alfredo Allen 208, 4200-135 Porto, Portugal; Cell Division and Cytoskeleton, Danish Cancer Society Research Center (DCRC), Strandboulevarden 49, 2100 Copenhagen, Denmark; UnIGENe, i3S - Instituto de Investigação e Inovação em Saúde, Universidade do Porto, Rua Alfredo Allen 208, 4200-135 Porto, Portugal; Instituto de Ciências Biomédicas Abel Salazar, Universidade do Porto, Rua Jorge Viterbo Ferreira, 228, 4050-313 Porto, Portugal; Cell Division Group, Experimental Biology Unit, Department of Biomedicine, Faculdade de Medicina, Universidade do Porto, Alameda Prof. Hernâni Monteiro, 4200-319 Porto, Portugal; Department of Cellular and Molecular Medicine, Faculty of Health Sciences, University of Copenhagen, Blegdamsvej 3B, 2200 Copenhagen, Denmark

## Abstract

Incorrect kinetochore-microtubule attachments during mitosis can lead to chromosomal instability, a hallmark of human cancers. Mitotic error correction relies on the kinesin-13 MCAK, a microtubule depolymerase whose activity in vitro is suppressed by α-tubulin detyrosination - a post-translational modification enriched on long-lived microtubules. However, whether and how MCAK activity required for mitotic error correction is regulated by microtubule tyrosination/detyrosination remains unknown. Here we found that microtubule detyrosination accumulates on correct, more stable, kinetochore-microtubule attachments, whereas constitutively high microtubule detyrosination near kinetochores compromised efficient error correction. Rescue experiments suggest that mitotic errors due to excessive microtubule detyrosination result from suppression of MCAK activity, without globally affecting kinetochore microtubule half-life. Importantly, MCAK centromeric activity was required and sufficient to rescue mitotic errors due to excessive microtubule detyrosination. Thus, microtubules are not just passive elements during mitotic error correction, and their tyrosination/detyrosination works as a ‘mitotic error code’ that allows centromeric MCAK to discriminate correct and incorrect kinetochore-microtubule attachments, thereby promoting mitotic fidelity.

## Introduction

Successful chromosome segregation during mitosis requires that each sister kinetochore is stably attached to microtubules oriented to opposite spindle poles (amphitelic attachments). However, due to the stochastic interactions between kinetochores and spindle microtubules early in mitosis, many chromosomes establish erroneous attachments that have been implicated in chromosomal instability (CIN), a hallmark of human cancers (Bakhoum and Cantley, 2018; Cimini et al., 2003). To prevent this, cells rely on error correction mechanisms that regulate microtubule dynamics at the kinetochore (Bakhoum et al., 2009). These mechanisms may act globally through the regulation of Cdk1 activity during early mitosis (Kabeche and Compton, 2013) or, more locally, by promoting microtubule detachment from kinetochores in response to low centromeric tension (Cimini et al., 2006; Liu et al., 2009). At the heart of this local error correction mechanism, the Aurora B kinase regulates the recruitment and/or activity of several centromeric/kinetochore proteins, including the kinesin-13 MCAK (Andrews et al., 2004; Bakhoum et al., 2009; Knowlton et al., 2006; Lan et al., 2004) - the most potent microtubule depolymerase present in animal cells (Desai et al., 1999). Therefore, mitotic error correction is currently viewed as a ‘blind’ process that results from the non-discriminatory renewal of microtubules at the kinetochore interface, regardless of their attachment status (i.e. correct or incorrect).

How MCAK mediates mitotic error correction has been difficult to determine, mostly due to its independent localization at centromeres/kinetochores and microtubule plus ends, and its global impact on spindle microtubule dynamics (Bakhoum et al., 2009; Domnitz et al., 2012; Huang et al., 2007; Kline-Smith et al., 2004; Rizk et al., 2009; Wordeman et al., 2007). Interestingly, the contribution of MCAK for kinetochore microtubule turnover appears to occur primarily in metaphase (Bakhoum et al., 2009), when most kinetochore-microtubule attachments are amphitelic and stabilized, and it does so without any measureable impact on microtubule polymerization dynamics associated with microtubule poleward flux (Ganem et al., 2005). This apparent paradox led us to hypothesize that centromeric MCAK is able to discriminate correct and incorrect kinetochore-microtubule attachments, independently of its global effect on spindle microtubule dynamics. In support of this hypothesis, active MCAK is enriched at centromeres/kinetochores of misaligned chromosomes (Andrews et al., 2004; Lan et al., 2004), as well as in aligned chromosomes with erroneous merotelic attachments (Knowlton et al., 2006).

In vitro reconstitution experiments have shown that MCAK activity is significantly suppressed by microtubule detyrosination (Peris et al., 2009; Sirajuddin et al., 2014), a post-translational modification on long-lived microtubules (Nieuwenhuis and Brummelkamp, 2019). Microtubule detyrosination has been recently implicated in mitosis and meiosis, neuronal processes and cognitive brain function, heart and skeletal mucle contraction, and cancer (Akera et al., 2017; Barisic et al., 2015; Chen et al., 2018; Erck et al., 2005; Kerr et al., 2015; Lafanechere et al., 1998; Liao et al., 2019; Pagnamenta et al., 2019; Robison et al., 2016). The detyrosination/tyrosination cycle involves the catalytic removal of the c-terminal tyrosine of most mammalian α-tubulin isoforms by tubulin carboxypeptidases (TCPs), such as the recently identified Vasohibin-SVBP complexes (Aillaud et al., 2017; Nieuwenhuis et al., 2017), followed by re-tyrosination of soluble α-tubulin by tubulin tyrosine ligase (TTL) (Ersfeld et al., 1993).

Here we combined powerful gene manipulation tools (including RNAi, small-molecule inhibition, protein overexpression and CRISPR-Cas9 gene editing) with state-of-the-art microscopy, including a novel super-resolution microscopy technique (Pereira et al., 2019), to investigate whether MCAK activity required for mitotic error correction is regulated by microtubule tyrosination/detyrosination. Our findings support a physiological role for microtubule tyrosination/detyrosination in the discrimination between correct and incorrect kinetochore-microtubule attachments, establishing a new paradigm in the control of mitotic fidelity.

## Results

### Microtubule detyrosination accumulates on correct, more stable, kinetochore-microtubule attachments

To investigate whether the mitotic error correction machinery ‘reads’ the tyrosination/detyrosination state of kinetochore microtubules we started by quantifying the ratio of detyrosinated/tyrosinated α-tubulin immediately adjacent to the kinetochore in distinct experimental conditions that favor a particular attachment configuration. These included short-lived syntelic (when both kinetochores from a pair are attached to microtubules from the same pole) or monotelic (when only one kinetochore from the pair is attached to microtubules) attachments, typically observed in monopolar spindles after kinesin-5 inhibition with monastrol (Kapoor et al., 2000); more stable amphitelic attachments that form during normal metaphase; and finally, hyperstabilized amphitelic attachments induced by treatment with 10 nM taxol for 1h. We found that correct amphitelic attachments show higher detyrosination near kinetochores when compared to incorrect/incomplete syntelic/monotelic attachments, and this tendency correlated with increased microtubule stability (Fig.1a-f). Thus, microtubule detyrosination near kinetochores increases with the establishment of correct, more stable, microtubule attachments.

**Figure 1.**
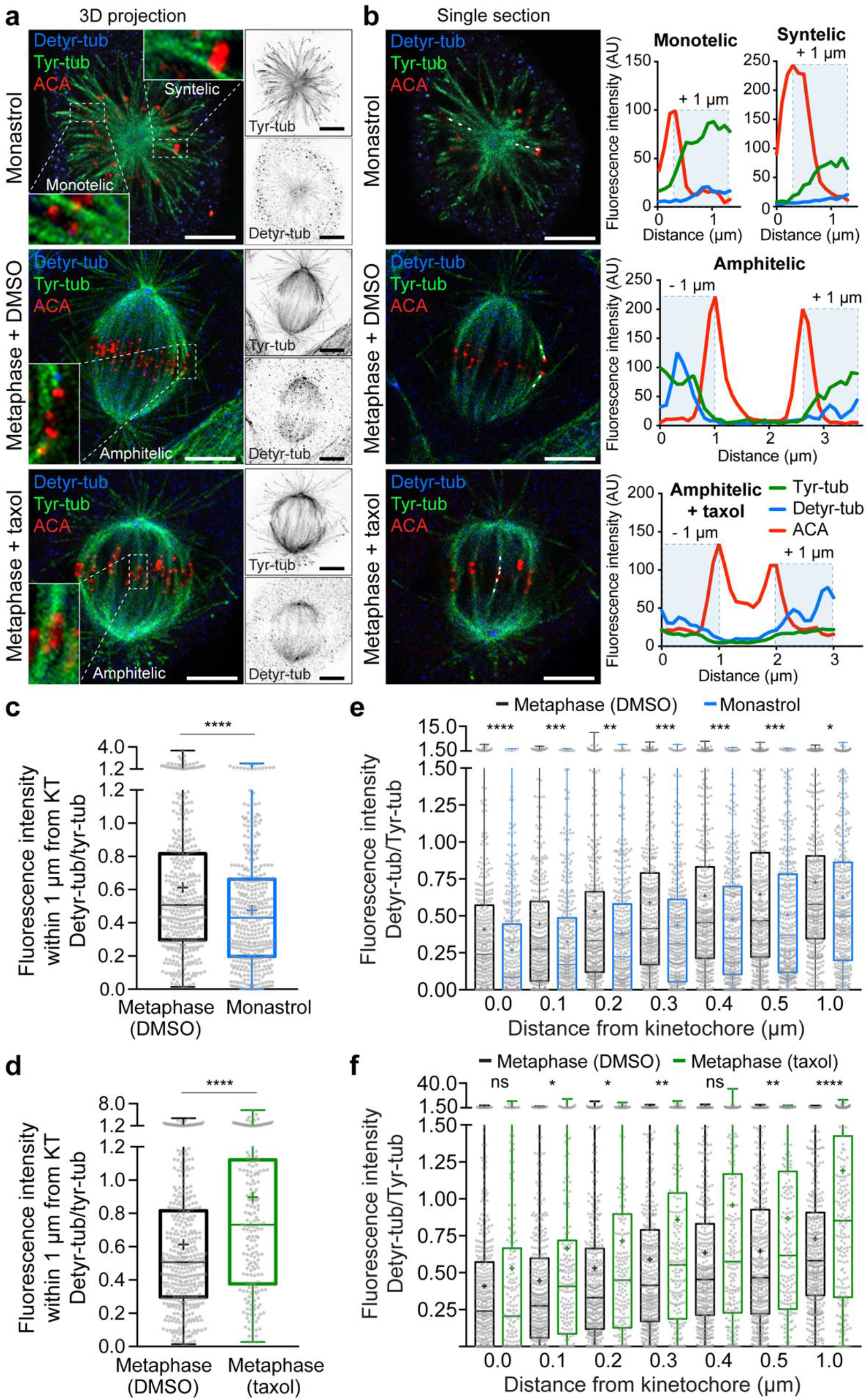
Microtubule detyrosination near kinetochores increases with the establishment of correct, more stable, microtubule attachments. **(a)** Confocal analysis of U2OS cells by immunofluorescence for detyrosinated and tyrosinated α-tubulin, and kinetochores with anti-centromere antibodies (ACA). Maximum intensity 3D projection of representative cells for each attachment configuration (zoomed images highlight monotelic/syntelic attachments - monastrol, amphitelic attachments - DMSO and hyper-stable amphitelic attachments - taxol). **(b)** Single sections from the respective confocal series shown in (a) with dashed lines indicating line-scanned representative kinetochore fibers (k-fibers). Scale bar, 5 μm. Line-scan plots of the selected kinetochore fibers indicated in (b). **(c, d)** Interquartile representation of the mean cumulative fluorescence intensity ratio between detyrosinated/tyrosinated α-tubulin within 1 μm distance from the kinetochore centroid in monotelic/syntelic attachments (monastrol: N=391 k-fibers, ~20 k-fibers/cell, 5 cells/experiment, 4 independent experiments) and hyper-stable amphitelic attachments (Taxol: N=199 k-fibers, ~20 k-fibers/cell, 5 cells/experiment, 2 independent experiments), respectively, compared to untreated amphitelic (DMSO: N=396 k-fibers, ~20 k-fibers/cell, 5 cells/experiment, 4 independent experiments) attachments. **(e, f)** Interquartile representation of the fluorescence intensity ratio between detyrosinated/tyrosinated α-tubulin as a function of distance from the kinetochore centroid from the same data set used in (c, d). Means are represented by “+”. p<0.001(****), p<0.005(***), p<0.01(**) and p<0.05(*), ns= non-significant; unpaired two-tailed t-test.

### Constitutively high microtubule detyrosination near kinetochores leads to mitotic errors

To directly investigate whether microtubule detyrosination impacts mitotic error correction we inactivated TTL either by RNAi or CRISPR/Cas9-mediated gene knockout (KO) in human U2OS cells. Both conditions caused a significant increase in the overall α-tubulin detyrosination levels, while reducing the tyrosinated α-tubulin pool (Supplementary Figs.1a and 2a). This was also the case on mitotic spindles, in which detyrosinated α-tubulin was increased along kinetochore fibers, including in the immediate vicinity of the kinetochore, and astral microtubules after TTL inactivation (Fig. 2a-c and Supplementary Figs. 1b, c and 2b). Live-cell imaging 72h after TTL RNAi in cells stably expressing H2B-histone-GFP/mCherry-α-tubulin revealed a consistent and significant increase in the frequency of anaphase cells with lagging chromosomes (Fig. 2d-f). Interestingly, this increase in chromosome missegregation gradually attenuated to control levels after chronic TTL inactivation (TTL KO), despite the continuous increase in detyrosinated α-tubulin over time (Supplementary Fig. 2c-f). This suggests the existence of compensatory or cellular adaptation mechanisms to the chronic loss of TTL, as shown previously for TTL KO mice (Erck et al., 2005). Importantly, experimental increase of α-tubulin detyrosination levels independently of TTL by overexpression of the TCP Vasohibin 1-SVBP led to an equivalent increase in anaphase cells with chromosome segregation errors when compared to TTL inactivation by RNAi (Fig. 3a, b, d, e). Conversely, RNAi-mediated depletion of Vasohibin 1 and 2 (with or without SVBP depletion), reduced chromosome missegregation events below control levels (Fig. 3c-e). Overall, these data indicate that experimental modulation of α-tubulin detyrosination levels on mitotic spindles, including in the vicinity of kinetochores, impacts chromosome segregation fidelity.

**Figure 2.**
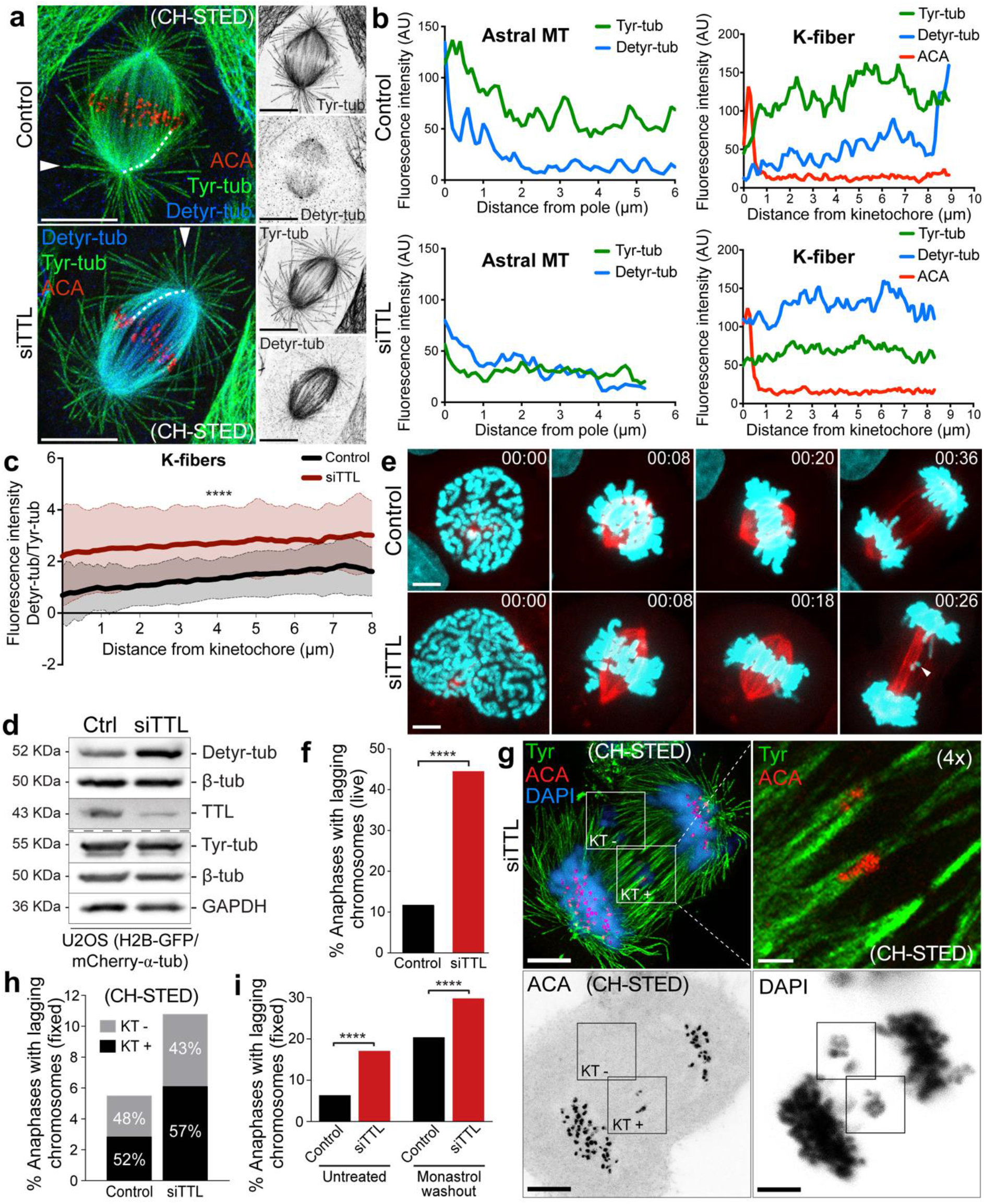
Constitutively high α-tubulin detyrosination in the vicinity of kinetochores impairs faithful chromosome segregation. **(a)** Confocal/CH-STED analysis of U2OS cells by immunofluorescence for detyrosinated tubulin (confocal), tyrosinated tubulin (CH-STED) and kinetochores/ACA (CH-STED). Dashed lines and arrowheads indicate representative line-scanned k-fibers and astral microtubules, respectively. Scale bars, 5 μm. **(b)** Graphic representation of the line-scans indicated in (a). **(c)** Quantification of the fluorescence intensity ratio between detyrosinated/tyrosinated α-tubulin along k-fibers (N=150 k-fibers; ~10 k-fibers/cell; 5 cells/experiment, 3 independent experiments for each condition). “0” corresponds to the kinetochore centroid. The mean values are represented by a thicker line and the standard deviation associated to each point is represented by a shaded band. **(d)** Protein lysates of U2OS cells stably expressing H2B-GFP/mCherry-α-tubulinm 72h after RNAi transfection were immunoblotted for detyrosinated and tyrosinated α-tubulin, TTL and β-tubulin. GAPDH was used as loading control. **(e)** Representative images from spinning-disk confocal time-series illustrating different stages of mitosis in control and TTL RNAi U2OS cells stably expressing H2B-GFP (cyan)/mCherry-α-tubulin (red). Pixels were saturated to allow the visualization of lagging chromosomes (arrowhead). Scale bar, 5 μm. Time is h:min. **(f)** Quantification of the percentage of anaphase cells with lagging chromosomes in control (11.7%) and TTL-deficient (siTTL) (44.3%) U2OS H2B-GFP/mCherry-α-tubulin cells [N> 60 cells, pool of 3 independent experiments; p<0.001 (****), logistic regression]. **(g)** Confocal/CH-STED analysis of a representative anaphase U2OS cell after siTTL by immunofluorescence for tyrosinated tubulin (CH-STED) and kinetochores/ACA (CH-STED). DNA was counterstained with DAPI (confocal). Inserts highlight kinetochore negative (KT-) and positive (KT+) lagging chromosomes. Scale bars, 5 μm. A 4X magnified maximum-projection (5 z-stacks) of the insert KT+ highlights the bi-orientation and stretching of a merotelic kinetochore. Scale bar, 1 μm. **(h)** Quantification of the relative percentage of KT- and KT+ lagging chromosomes in control and TTL RNAi cells (N>1000 anaphase cells, pool of 2 independent experiments, 3 replicates per experiment). **(i)** Quantification of the percentage of anaphase cells with lagging chromosomes in control and TTL-depleted (siTTL) cells in an asynchronous (untreated) cell population (control: N=566 anaphase cells; siTTL: N=386 anaphase cells, pool of 3 independent experiments, p<0.001(****), logistic regression) and upon monastrol washout (control: N=3977 anaphase cells; siTTL: N=3130 anaphase cells, pool of 11 independent experiments, p<0.001(****), logistic regression).

**Figure 3.**
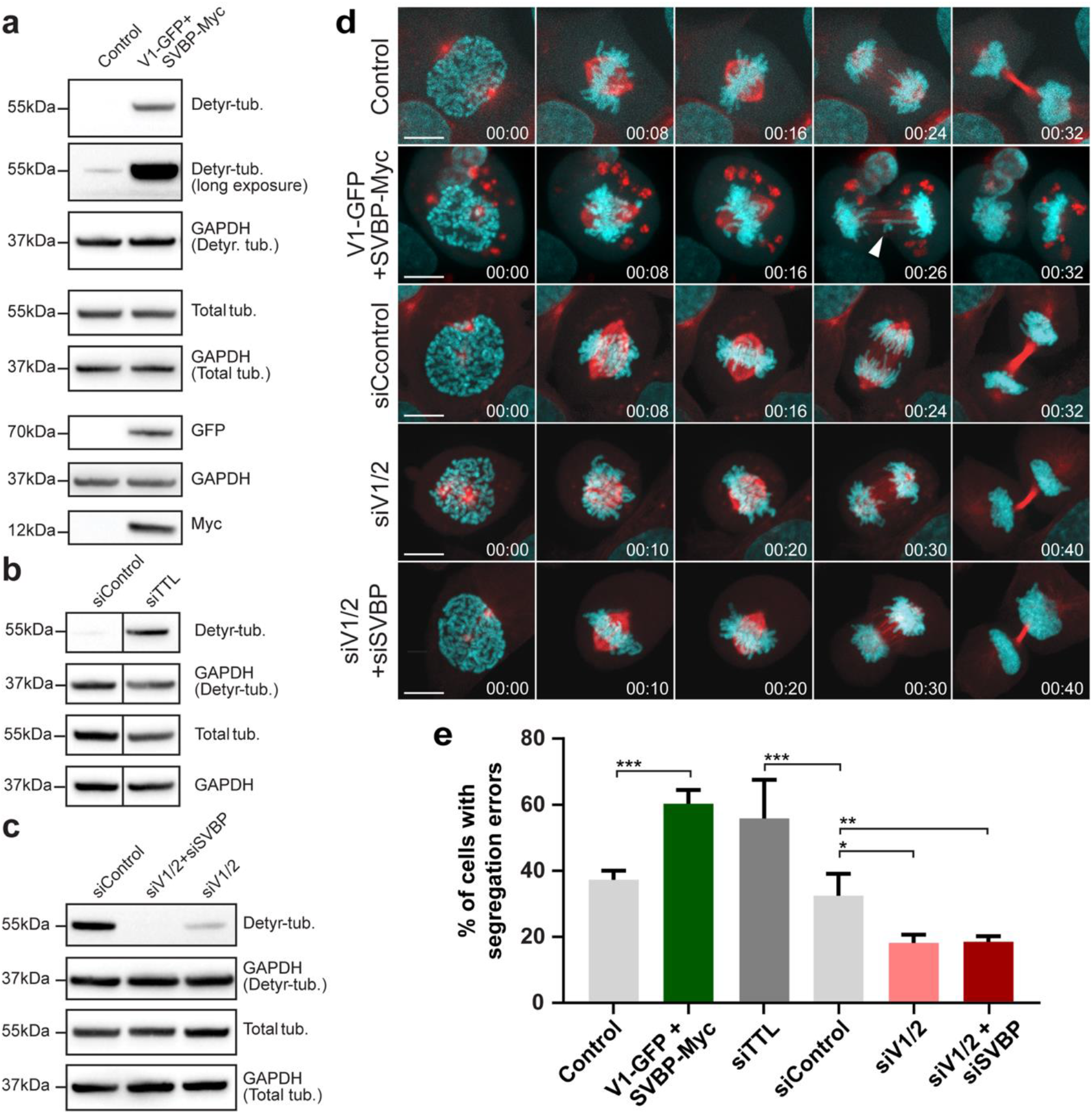
Modulation of α-tubulin detyrosination levels by manipulation of Vasohibins-SVBP impacts chromosome segregation fidelity. (**a-c**) Immunoblot analysis of detyrosinated α-tubulin and total α-tubulin levels after VASH1-SVBP overexpression, RNAi mediated knockdown of TTL or VASH1 and VASH2 (with/without SVBP), respectively. GAPDH was used as loading control. The expression of VASH1-GFP and SVBP-myc was observed using anti-GFP and anti-myc antibodies, respectively. (**d**) Representative images from spinning-disk confocal time-series displaying different stages of mitosis in U2OS cells stably expressing H2B-GFP (cyan)/mCherry-α-tubulin (red) transfected with the indicated plasmids and siRNAs. White arrowhead points to a segregation error during anaphase. Scale bar, 10 μm. Time in h:min. (**e**) Quantification of the percentage of anaphase cells with segregation defects (lagging chromosomes and DNA bridges not discriminated) from spinning-disk confocal imaging of U2OS cells stably expressing H2B-GFP/mCherry-α-tubulin transfected with the indicated plasmids and siRNAs. N(number of cells, number of independent experiments): Control(GFP)(38, 4); VASH1-GFP+SVBP-myc (38, 4); siTTL (28, 2); siControl (121, 5); siVASH1/2 (91, 4) and siVASH1/2+siSVBP (45, 3). p<0.05(*), p<0.01(**), one-way ANOVA.

### Mitotic errors due to excessive microtubule detyrosination result from incorrect kinetochore-microtubule attachments

Persistent merotelic attachments in which a single kinetochore is attached to microtubules from both spindle poles are the main source of anaphase lagging chromosomes in mammalian cells (Cimini et al., 2003). To investigate whether the lagging chromosomes observed after experimental increase in α-tubulin detyrosination result from the formation of merotelic attachments, we used super-resolution CH-STED microscopy (Pereira et al., 2019) in fixed cells after TTL RNAi. Relative to conventional 2D-STED, this novel sub-diffraction technique improves contrast in highly complex environments, such as the mitotic spindle. Accordingly, we observed a significant increase in lagging chromosomes with single stretched kinetochores attached to microtubules from both poles, confirming the formation of merotelic attachments (Fig. 2g, h). Noteworthy, TTL-depleted cells also showed a significant increase in acentric (i.e. without kinetochores) chromosome fragments, probably resulting from the sheer stress undergone by the centromere during correction of merotelic attachments during anaphase (Cimini et al., 2003; Guerrero et al., 2010) or due to chromosome shattering of missegregated chromosomes in the previous division cycle (Crasta et al., 2012). Taken together, these data indicate that a significant fraction of the lagging chromosomes observed after experimental increase of α-tubulin detyrosination derive from erroneous merotelic attachments.

### Excessive microtubule detyrosination impairs efficient error correction during mitosis

To distinguish whether increased microtubule detyrosination increases error formation and/or prevents efficient error correction, we induced the assembly of monopolar spindles with monastrol, which favors the formation of merotelic attachments during spindle bipolarization after monastrol washout (Cimini et al., 2003; Kapoor et al., 2000). This treatment led to an equivalent increase in lagging chromosomes in both control and TTL-depleted anaphase cells (Fig. 2i), suggesting that the increased frequency of mitotic errors observed after experimental increase of α-tubulin detyrosination were not due to an increased error formation rate. This conclusion was substantiated by the observation that the timing of centrosome separation at nuclear envelope breakdown, an important variable that has been linked to increased error formation during mitosis (Silkworth et al., 2012), was indistinguishable between control and TTL-depleted cells (Supplementary Fig. 3a-c). Thus, increased microtubule detyrosination impairs efficient error correction during mitosis.

### Excessive microtubule detyrosination does not interfere with global kinetochore-microtubule dynamics

Given that microtubule detyrosination correlates with microtubule stability, which has been proposed as the main cause of persistent merotelic attachments behind CIN (Bakhoum et al., 2009; Cimini et al., 2003; Cimini et al., 2006), we investigated whether constitutive amplification of microtubule detyrosination affects global kinetochore microtubule stability. To do so, we used fluorescence dissipation after photoactivation (FDAPA) of GFP-α-tubulin to measure kinetochore microtubule turnover in control and TTL-depleted cells, as well as in MCAK-depleted cells used as positive control (Bakhoum et al., 2009; Ferreira et al., 2018). The resulting decay curve fits a double exponential (R^2^>0.99), reflecting the presence of two microtubule populations: a fast fluorescence decay population that has been attributed to non-kinetochore microtubules; and a slow fluorescence decay population thought to correspond to more stable kinetochore microtubules (Fig. 4a, b) (Zhai et al., 1995). From the fit we extracted the relative percentages of stable versus unstable microtubules, and their respective half-lives (t1/2). We found that kinetochore microtubule turnover increased from prometaphase to metaphase in all conditions, but was indistinguishable between control and TTL-depleted cells at each given stage (Fig. 4c). In line with these data, measurement of inter-kinetochore distances as a proxy for tension on amphitelic attachments was also indistinguishable between control and TTL-depleted cells (Supplementary Fig. 1d). In contrast, MCAK-depleted cells had slightly more stable kinetochore microtubules specifically during metaphase (Fig. 4c), as shown previously (Bakhoum et al., 2009). Altogether, these data are consistent with previous measurements of microtubule half-life after experimental increase of microtubule detyrosination in interphase (Webster et al., 1990), and demonstrate that microtubule detyrosination does not interfere with global kinetochore-microtubule dynamics.

**Figure 4.**
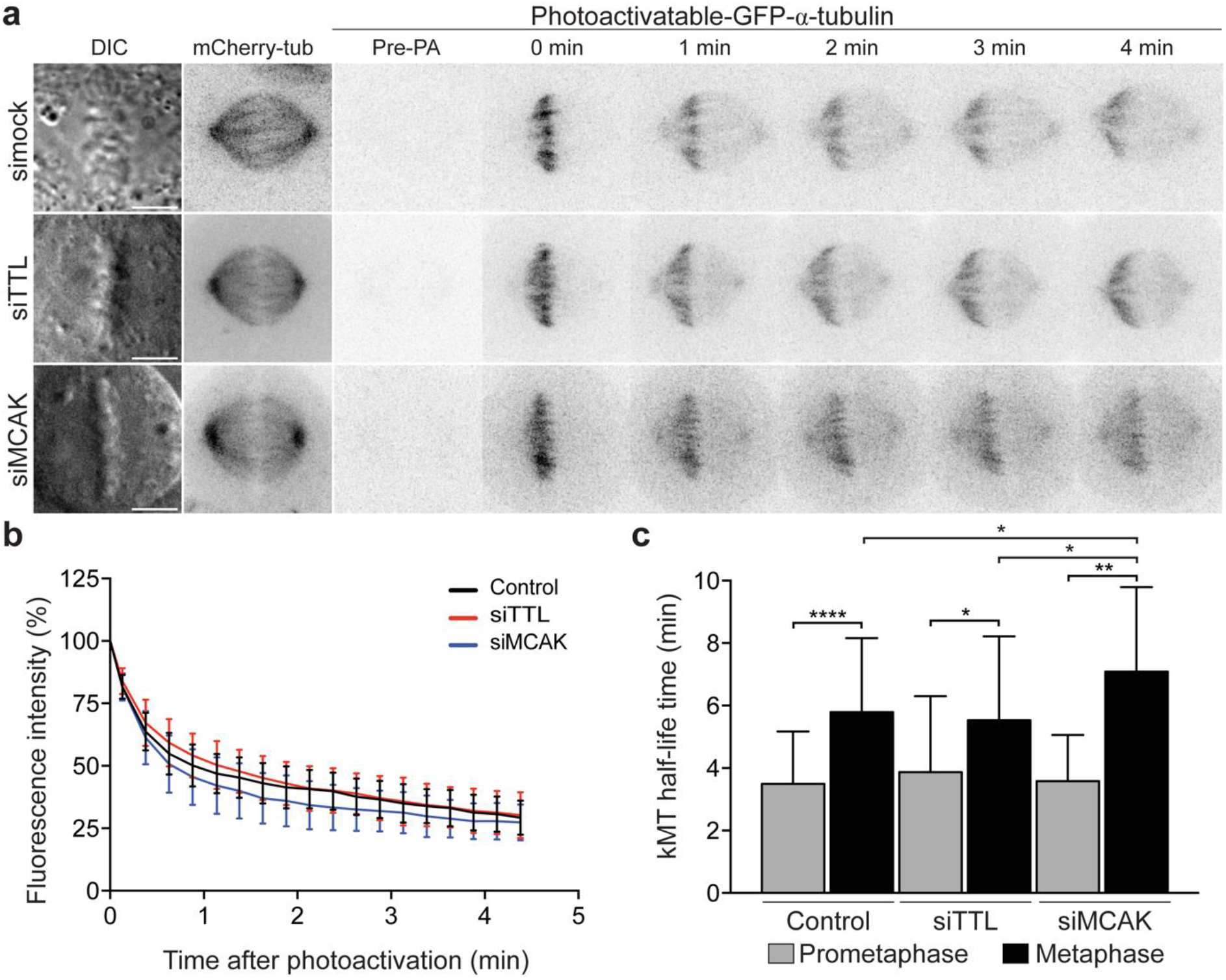
Microtubule detyrosination does not interfere with global kinetochore-microtubule dynamics. **(a)** Representative images of differential interference contrast (DIC) and time-lapse fluorescent images of metaphase spindles in control, TTL-deficient (siTTL) and MCAK-deficient (siMCAK) U2OS cells stably expressing Photoactivatable-GFP/mCherry-tubulin before photoactivation (Pre-PA) and at the indicated time points (min) after photoactivation of GFP-tubulin fluorescence. Scale bar, 5 μm. **(b)** Normalized fluorescence intensity after photoactivation in all conditions for metaphase cells. Data points represent mean ± standard deviation. **(c)** Calculated kinetochore microtubule (kMT) half-life under different conditions (control: N=32 cells, pool of 6 independent experiments; siTTL: N=17 cells, pool of 3 independent experiments; siMCAK: N=30 cells, pool of 2 independent experiments, p<0.05(*), p<0.01(**), p<0.001(****), unpaired two-tailed t-test). Bars indicate mean values/cell and error bars represent standard deviation.

### Impaired error correction due to excessive microtubule detyrosination results from compromised MCAK activity

Interestingly, while TTL or MCAK depletion significantly increased the frequency of anaphase cells with lagging chromosomes, TTL-depleted cells showed a clear enrichment of detyrosinated α-tubulin along kinetochore microtubules, whereas MCAK-depleted cells strongly accumulated detyrosinated α-tubulin preferentially on astral microtubules (Fig. 5a-c). These striking differences in the distribution of detyrosinated α-tubulin imply that global perturbation of the two proteins impacts kinetochore-microtubule attachments through different mechanisms. Indeed, MCAK was proposed to control the formation of robust kinetochore-microtubule attachments both by regulating the length of non-kinetochore microtubules through a role at microtubule plus ends (Domnitz et al., 2012), and by controlling chromosome directional switching by a specific role at centromeres (Kline-Smith et al., 2004; Wordeman et al., 2007). Additionally, microtubule detyrosination was also recently shown to regulate MCAK activity required for normal astral microtubule length (Liao et al., 2019). To address whether and how microtubule detyrosination impairs MCAK activity required for mitotic error correction, we set up rescue experiments either by overexpressing exogenous full-length EGFP-MCAK (MCAKFL) or by enhancing endogenous MCAK activity with the small-molecule agonist UMK57 (Orr et al., 2016) in TTL-deficient cells. We found that both treatments partially rescued the frequency of anaphase cells with lagging chromosomes observed after TTL depletion (Fig. 4a-d), suggesting that the impaired error correction due to increased microtubule detyrosination results from compromised MCAK activity.

**Figure 5.**
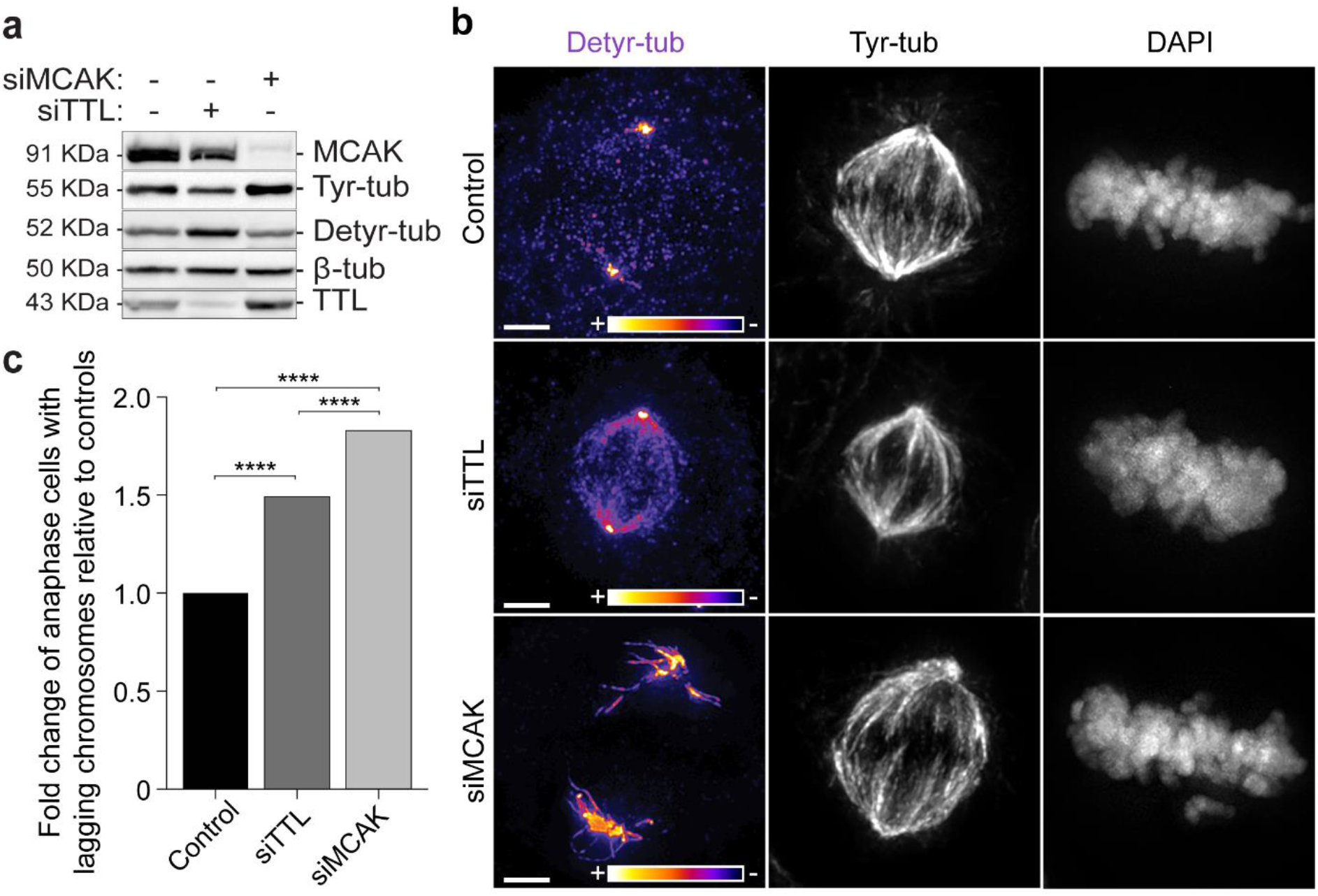
Distinct distribution of detyrosinated α-tubulin after TTL or MCAK depletion. (**a**) Protein lysates from parental U2OS cells 72h after RNAi were immunoblotted for MCAK, Tyr-tub, Detyr-tub, β-tubulin and TTL. β-tubulin was used as loading control. (**b**) 3D-deconvolution analysis of U2OS cells after immunofluorescence for Tyr-tub and Detyr-tub. DNA was counterstained with DAPI. The fire look-up table was used to map fluorescence intensity of Detyr-tub. Scale bar, 5 μm. (**c**) Quantification of the fold change of anaphase cells with lagging chromosomes after monastrol washout for each condition relative to controls [Control: N= 877 anaphase cells, siTTL: N= 1142 anaphase cells, siMCAK: N= 743 anaphase cells, pool of 3 independent experiments for each condition; p<0.001(****), logistic regression].

### Centromeric MCAK activity is required and sufficient to rescue error correction in response to a constitutive increase of microtubule detyrosination, without globally changing kinetochore microtubule dynamics

Additional rescue experiments with MCAK mutant constructs that compromise microtubule depolymerizing activity (MCAKhypir) or centromeric localization (ΔNMCAK) (Domnitz et al., 2012; Wordeman et al., 2007) further suggest that microtubule detyrosination impairs error correction by specifically inhibiting MCAK activity at centromeres (Fig. 6a-c, e). To directly test this possibility, we constitutively targeted a GFP-tagged minimal motor domain of MCAK to the centromere fused with the centromere binding domain of CENP-B, with (GCPBM) or without (GCPBMhypir) microtubule depolymerising activity (Wordeman et al., 2007). We found that overexpression of either GCPBM or GCPBMhypir did not cause any measurable differences in mitotic spindle and astral microtubule length (Fig. 6a-c, f, g). Yet, GCPBM, but not its inactive counterpart, was sufficient to significantly rescue the abnormally high frequency of anaphase cells with lagging chromosomes observed in TTL-depleted cells (Fig. 6e), suggesting that, in addition to affecting MCAK’s role on astral microtubules (Liao et al., 2019), microtubule detyrosination also inhibits a restricted pool of centromeric MCAK required for error correction. To directly test this, we used FDAPA in U2OS cells stably expressing photoconvertible mEos-α-tubulin to measure global kinetochore microtubule half-life after overexpression of GCPBM. We found that overexpression of GCPBM abolished the stabilization of kinetochore microtubules typically observed from prometaphase to metaphase, without significantly affecting overall kinetochore microtubule half-life (Fig. 7a-c). Thus, MCAK depolymerizing activity at centromeres is required and sufficient to rescue error correction in response to a constitutive increase of microtubule detyrosination, independently from its global role in the regulation of spindle microtubule dynamics.

**Figure 6.**
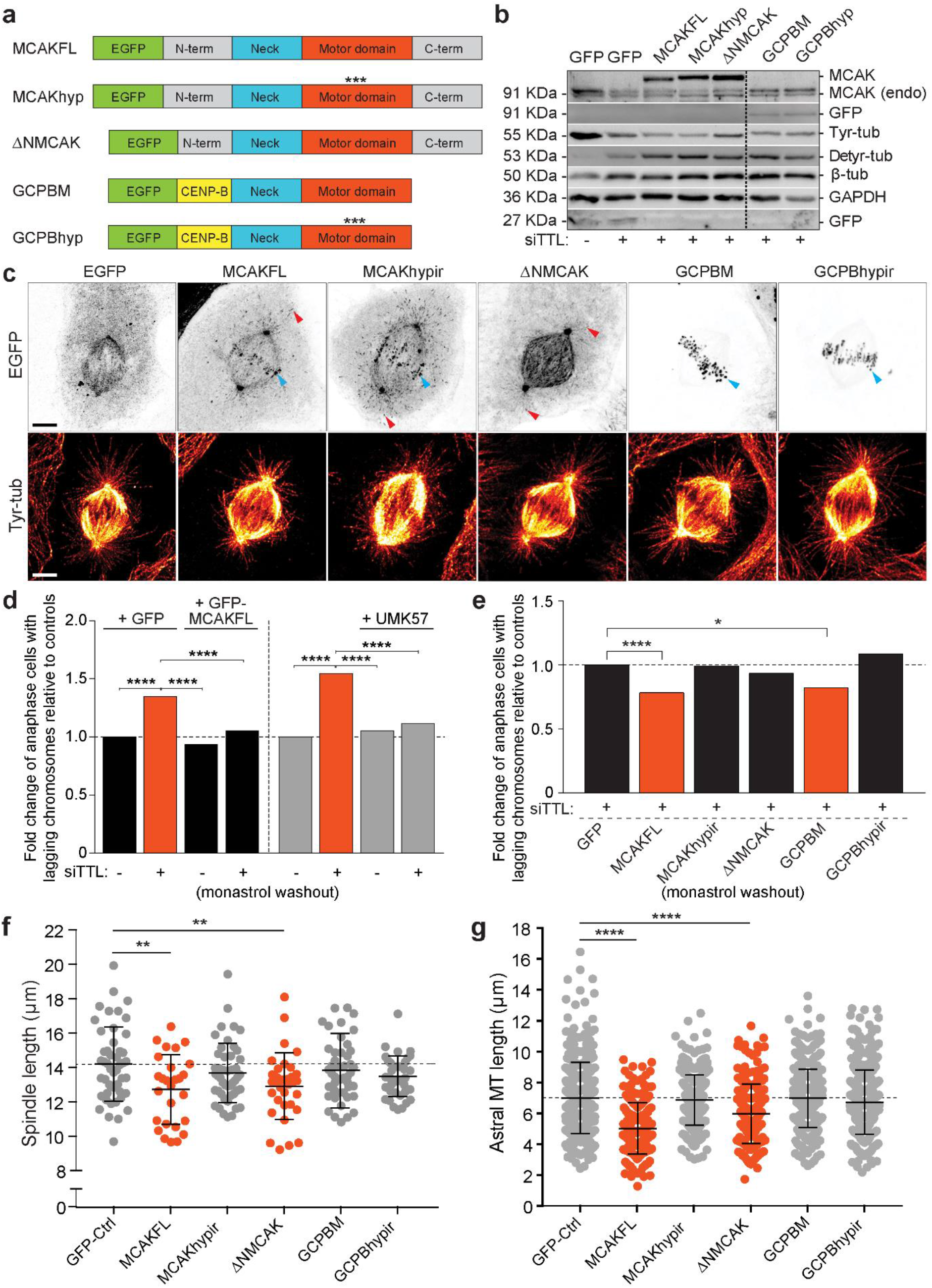
MCAK depolymerizing activity at centromeres is required and sufficient to rescue error correction in response to a constitutive increase of microtubule detyrosination. (**a**) Diagram of the MCAK constructs overexpressed in this study. (**b**) Protein lysates of U2OS cells 24-36 h after DNA transfection with the different MCAK constructs after TTL depletion were immunoblotted for GFP, MCAK, Tyr-tub, Detyr-tub and β-tubulin. GAPDH was used as loading control. **(c)** 3D-deconvolution analysis of U2OS cells overexpressing different EGFP-MCAK constructs and immunostained for tyrosinated α-tubulin (red-hot look-up table). Scale bars, 5 μm. **(d)** Quantification of the fold change of anaphase cells with lagging chromosomes relative to controls in rescue experiments using MCAKFL overexpression (N>1500 anaphase cells, pool of 5 independent experiments, p<0.001(****), logistic regression) and MCAK pharmacological enhancement with UMK57 (N>1000 anaphase cells, pool of 4 independent experiments, p<0.001(****), logistic regression). **(e)** Quantification of the fold change of anaphase cells with lagging chromosomes relative to controls in rescue experiments using different EGFP-MCAK constructs. Significant rescue was observed upon overexpression of MCAKFL (N>1500 anaphase cells, pool of 5 independent experiments, p<0.001(****), logistic regression) and GCPBM (N>900 anaphase cells, p=0.011(*), pool of 4 independent experiments, logistic regression). **(f)** Quantification of the metaphase spindle length 24-48h after transfection with different EGFP-MCAK constructs [GFP-Control: N=47 cells, MCAKFL: N=27 cells, MCAKhypir: N=47 cells, ΔNMCAK: N=33 cells, GCPBM: N=46 cells, GBPBMhypir: N=38 cells, pool of 3 independent experiments for each condition, except for GCPBMhypir (pool of 2 independent experiments), p<0.01 (**), unpaired two-tailed t-test]. **(g)** Quantification of the astral microtubule (MT) length [~10 astral MTs/cell; GFP-Control: N=471 astral MTs, MCAKFL: N=264 astral MTs, MCAKhypir: N=470 astral MTs, ΔNMCAK: N=326 astral MTs, GCPBM: N=456 astral MTs, GBPBMhypir: N=378 astral MTs, pool of 3 independent experiments for each condition, except for GCPBMhypir (pool of 2 independent experiments), p<0.001 (****), unpaired two-tailed t-test].

**Figure 7.**
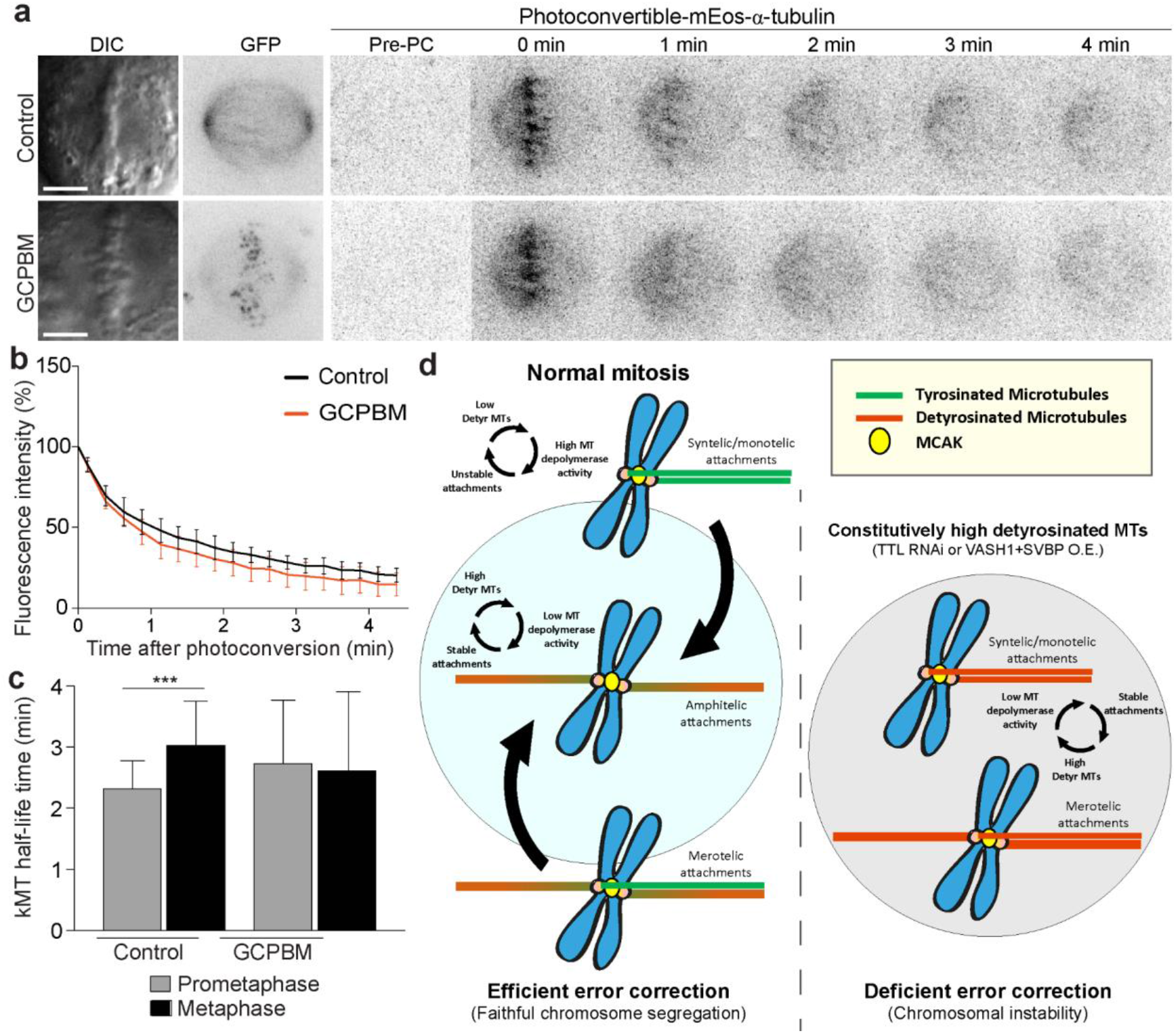
Centromeric MCAK does not significantly alter global kinetochore microtubule dynamics. **(a)** Representative images of DIC and time-lapse fluorescent images of metaphase spindles of U2OS cells stably expressing mEos-α-tubulin. Photoconversion was performed after overexpression of GFP (control) and GCPBM. Images before photoconvertion (Pre-PC) and at the indicated times points (min) after photoconversion are displayed. Scale bar, 5 μm. (**b**) Normalized fluorescence intensity after photoconversion of mEos-α-tubulin in U2OS cells in metaphase (GFP-Control: N=9 cells from 5 independent experiments and GCPBM: N=6 cells from 3 independent experiments). Data points represent mean ± standard deviation. (**c)**Calculated kMT half-life in control and GCPBM overexpressing cells in prometaphase (GFP-Control: N=21 cells, and GCPBM: N=25 cells, pool of 5 independent experiments, p=0.0847, unpaired two-tailed t-test) and metaphase (GFP-Control: N=9 cells from 5 independent experiments and GCPBM: N=6 cells from 3 independent experiments, p=0.4351, unpaired two-tailed t-test) using the fluorescence decay after photoconversion. Bars indicate mean values/cell and error bars represent standard deviation. (**d**) Proposed model for the discrimination of mitotic errors by microtubule detyrosination (see main text for description).

## Discussion

The prevalent view is that microtubules are ‘passive’ elements during mitotic error correction. The results reported in this work support a model in which differential microtubule tyrosination/detyrosination near the kinetochore works as a ‘mitotic error code’ that allows microtubule depolymerizing enzymes located at centromeres/kinetochores to discriminate between correct and incorrect kinetochore-microtubule attachments. According to this model (Fig. 7d), initial attachments (including incorrect ones) established during early mitosis are essentially formed by tyrosinated microtubules, and thus permissive to depolymerize and detach. As chromosomes bi-orient and centromeric tension develops, amphitelic attachments become more stabilized and consequently more detyrosinated, leading to their further stabilization. While allowing the selective preservation of correct amphitelic attachments, this microtubule-based feedback mechanism explains how de novo merotelic attachments on already bi-oriented chromosomes can be surgically prevented/corrected, given that the low detyrosination of newly attached microtubules would favor their destabilization. Thus, spindle microtubules are not just passive elements during mitotic error correction, and their detyrosination/tyrosination works as an active signalling mechanism that discriminates mitotic errors to promote mitotic fidelity. These findings have implications for the use of microtubule stabilizing drugs, such as taxol and its derivatives, in the treatment of human cancers, many of which acquire resistance by modulating MCAK activity (Ganguly et al., 2011a; Ganguly et al., 2011b).

## Acknowledgements

We would like to thank Linda Wordeman for kindly sharing MCAK constructs, Ben Kwok and Duane Compton for the UMK57, Stephan Geley and René Medema for the U2OS lines, Jakob Nilsson for gateway plasmids, Martina Barisic for technical support and the Biosciences Screening Scientific platform at i3S. This work was funded by the European Research Council (ERC) under the European Union’s Horizon 2020 research and innovation programme (grant agreement No 681443) and FLAD Life Science 2020 (to H.M.), and by grants from the Danish Cancer Society Scientific Committee (KBVU; R146-A9322) and the Lundbeck Foundation (R215-2015-4081 (to M.B). L.T.F. was supported by a studentship from Fundação para a Ciência e a Tecnologia (SFRH/BD/79174/2011).

## Author contributions

Investigation, formal analysis and validation (L.T.F., B.O., G. R., C.G.B., J.T.L.); conceptualization, supervision, and funding acquisition (H.M., M.B., J.G.F.); resources, methodology and software (A.J.P.); validation and formal analysis (C.L.); writing - original draft (L.T.F., H.M.); writing – review and editing (L.T.F., G. R., C.G.B., J.T.L., H.M., M.B.); project administration (H.M.).

## Conflict of interests

The authors declare no conflict of interest.

## Materials and Methods

### Cell lines and plasmids

All cell lines were cultured at 37°C in 5% CO_2_ atmosphere in Dulbecco’s modified medium (DMEM, Gibco™, Thermofisher) containing 10% fetal bovine serum (FBS, Gibco™, Thermofisher). Parental, mEos-α-tubulin and H2B-GFP/mCherry-α-tubulin U2OS cells were kindly provided by S. Geley (Innsbruck Medical University, Innsbruck, Austria), whereas the PA-GFP-α-tubulin/mCherry-α-tubulin U2OS cells were kindly provided by R. Medema (NKI, Amsterdam, The Netherlands). Stable H2B-GFP/mRFP-α-tubulin U2OS cells were generated in this study by lentiviral transduction.

### Constructs and transfections

To express fluorescently-tagged proteins, cells were transfected using Lipofectamine™ 2000 (Thermofisher) with 2 μg of each constructs. MCAK constructs (kindly provided by L. Wordeman, University of Washington, USA): EGFP-MCAK full-length, EGFP-MCAKhypir, EGFP-ΔN-MCAK, GCPBM and GCPBMhypir(Domnitz et al., 2012; Wordeman et al., 2007); Human VASH1 (NM_014909.5) and SVBP (NM_199342.4) encoding gateway-compatible vectors (kind gift from J. Nilsson, Center for Protein Research, Copenhagen, Denmark) were used to create expression plasmids with C-terminal eGFP and 3X-myc tags by LR recombination (Invitrogen) according to manufacturers instructions. Plasmid encoding eGFP alone was used as control. Expression of VASH1 and SVBP were performed as described before(Liao et al., 2019). Briefly, cells were transfected with 1 μg of the two plasmids using GeneJuice (Merck) transfection reagent according to manufacturer’s protocol. To perform RNA interference (RNAi)-mediated protein depletion, cells were plated at 30-50% confluence onto 22 × 22 mm No. 1.5 glass coverslips and cultured for 1-2 h in DMEM supplemented with 5% of FBS before transfection. RNAi transfection was performed using Lipofectamine™ RNAiMAX reagent (Thermofisher) with 50 nM of validated siRNA oligonucleotides against human TTL 5’-GUGCACGUGAUCCAGAAAU-3’, MCAK 5’-GAUCCAACGCAGUAAUGGU-3’, VASH1 5’-CCAGACUAGGAUGCUUCUG-3’, VASH2 5’-GGAUAACCGUAACUGAAGU-3’, SVBP 5’-GCAGAGAUCUAUGCCCUCA-3’ and control (scramble) siRNA 5’-UGGUUUACAUGUCGACUAA-3’, diluted in serum-free medium (Opti-MEM™, Thermofisher). Mock transfected or scramble siRNA results were indistinguishable. The TTL knockout cells were generated by CRISPR/Cas9-mediated genome editing, using a lentiviral backbone containing both the *Streptococcus pyogenes* Cas9 (*sp*Cas9) nuclease and the single guide RNA (sgRNA) scaffold (lentiCRISPRv2). Two 20 bp sgRNA (sgTTL, 5’-CACCGAACAGCAGCGTCTACGCCG-3’ and sgCtrl, 5’-AAACCGGCGTAGACGCTGCTGTTC-3’) (Sigma-Aldrich) were designed to target TTL gene and cloned into the lentiviral vector pLenti-CRISPR-v2 (#52961, addgene). TTL knockout cells were selected by their resistance to puromycin (2 μg/mL, Calbiochem) and confirmed by western blot analysis.

### Drug treatments

To induce lagging chromosomes by inhibition of kinesin-5 we added 100 μM Monastrol (Tocris, Bioscience) 12-16 h before washout. Microtubule stabilization was obtained using 10 nM Taxol for 1 h (Sigma-Aldrich). Mitotic arrest at metaphase was obtained using 10 μM MG132 (Calbiochem). Live-cell analysis using MG132 was performed in less than 2 h in the presence of the drug to avoid cohesion fatigue. MCAK activity was enhanced using 250 μM of UMK57(Orr et al., 2016) for 12-16 h before fixation. An equivalent volume of DMSO was used as control for each drug treatment.

### Immunofluorescence

U2OS cells were fixed with cold methanol (−20°C) for 4 min or 0.1-0.25% Gluteraldehyde + 4% Paraformaldehyde (Electron Microscopy Sciences) for 10 min. Autofluorescence was quenched by 0.1% sodium borohydride (Sigma-Aldrich) after aldehyde fixation and cells permeabilized with 0.5% Triton X-100 (Sigma-Aldrich) for another 10 min. Tyrosinated α-tubulin and detyrosinated α-tubulin were immunostained using rat monoclonal anti-tyrosinated α-tubulin clone YL1/2 (1:100-500, Bio-Rad) and rabbit polyclonal anti-detyrosinated α-tubulin (1:250-2000)(Liao et al., 2019), respectively. Other primary antibodies used were human anti-centromere antibodies (ACA, 1:100-2000, kind gift from B. Earnshaw, Welcome Trust Centre for Cell Biology, University of Edinburgh, UK, or S. Geley). GFP-tagged constructs were visualized by means of direct fluorescence. Alexa Fluor 488 (1:200-2000, Themofisher), STAR-580 and STAR-RED (1:200, Abberior) were used as secondary antibodies and DNA was counterstained with 1 μg/mL DAPI (4’,6’-diamino-2-fenil-indol, Sigma-Aldrich).

### Image Acquisition

3D wide-field images were acquired using an AxioImager Z1 (100× Plan-Apochromatic oil differential interference contrast objective lens, 1.46 NA, ZEISS) equipped with a CCD camera (ORCA-R2, Hamamatsu) operated by Zen software (ZEISS). Blind deconvolution of 3D image data sets was performed using Autoquant X software (Media Cybernetics). Confocal and super-resolution CH-STED images were acquired using Abberior Instruments ‘Expert Line’ gated-STED coupled to a Nikon Ti microscope. An oil-immersion 60× 1.4 NA Plan-Apo objective (Nikon, Lambda Series) and pinhole size of 0.8 Airy units was used in all confocal acquisitions. Super-resolution images were acquired using a coherent hybrid (CH−) STED beam(Pereira et al., 2019).

### Quantification of the fluorescence intensity along astral microtubules and kinetochore fibers

Detyrosination/tyrosination profile along astral microtubules and kinetochore fibers (including in a sub-region 1 μm distant from the kinetochore was performed using ImageJ by drawing a line segment along the region of interest followed by plotting the fluorescence intensity values (command plot profile at each channel) as a function of distance from the kinetochore centroid determined by ACA signal. Background was quantified in a region outside the spindle and subtracted to each channel.

### Quantification of detyrosination at the kinetochores

The fluorescence intensity signal of tubulin detyrosination was measured directly at kinetochores in a 9×9 pixel square region of interest (ROI) using LAPSO (MATLAB) and normalized to the intensity signal of ACA in the same ROI. Background fluorescence was measured outside the ROI and subtracted to each kinetochore. The mean values of all kinetochores quantified in one cell were plotted in a scattered dot plot.

### Quantification of spindle and astral microtubule length

The spindle length was calculated as the distance between the two centrosome position adjusted to the focal plane according to the equation: Spindle length = √[d(centrosome1−centrosome2, xy)^2^ + d(centrosome1−centrosome2, z)^2^, d=distance. Astral microtubule length was measured as the distance between the centrosome and the microtubule distal tip using maximum-projection images.

### Quantification of the inter-kinetochore distance

Inter-kinetochore distances were measured using a custom program written in MATLAB 8.1 (QUANTA), which determines the 3D-distance between the centroid of the sister kinetochores.

### Time-lapse microscopy

For phenotypic analysis of TTL depletion (siTTL), U2OS H2B-GFP/mCherry-α-tubulin cells were cultured in glass coverslips and assembled into 35 mm magnetic chambers (14 mm, No. 1.5, MatTek Corporation). Cell culture medium was replaced with phenol-red-free DMEM CO_2_-independent media (Invitrogen) supplemented with 10% FBS. Time-lapse imaging was performed in a heated chamber (37 °C) using a 100× 1.4 NA Plan- Apochromatic differential interference contrast objective mounted on an inverted microscope (TE2000U; Nikon) equipped with a CSU-X1 spinning-disk confocal head (Yokogawa Corporation of America) and with two laser lines (488 nm and 561 nm). Images were acquired with an iXon+ EM-CCD camera (Andor Technology). Eleven 1 μm-separated z-planes covering the entire volume of the mitotic spindle were collected every 2 min. The phenotypic analysis of U2OS H2B-GFP/mCherry-α-tubulin cells overexpressing GFP and VASH1-GFP+SVBP-Myc, as well as the analysis of VASH1/2, VASH1/2+SVBP and additional TTL depletion (used as positive control) in the same experimental data set, was performed using a Plan-Apochromat DIC 63x/1.4NA oil objective mounted on an inverted Zeiss Axio Observer Z1 microscope (Marianas Imaging Workstation from Intelligent Imaging and Innovations Inc. (3i), Denver, CO, USA), equipped with a CSU-X1 spinning-disk confocal head (Yokogawa Corporation of America), iXon Ultra 888 EM-CCD camera (Andor Technology) and four laser lines (405 nm, 488 nm, 561 nm and 640 nm). Fifteen 1 μm thick z-planes were collected every 2 min for 2-3 h. Image processing was performed in ImageJ. All displayed images represent maximum-intensity projections of z-stacks. Additional phenotypic live-imaging analysis of TTL KO H2B-GFP/mCherry-α-tubulin cells and respective controls was performed in an IN Cell Analyzer 2000 microscope (GE Healthcare), equipped with temperature and CO_2_ controller, using a Nikon 40x/0.95 NA Plan Fluor objective and a large chip CCD Camera (CoolSNAP K4) with a pixel array of 2048×2048. Single planes were acquired every 2 min.

### Error correction assay

All U2OS cell lines used in this assay were seeded onto 22 × 22 mm No. 1.5 glass converslips and cultured at 37°C in 5% CO_2_ atmosphere in DMEM (Gibco™, Thermofisher) supplemented with 10% FBS (Gibco™, Thermofisher) for 72 h. 100 μM of Monastrol (Tocris, Bioscience) was added 12-16 h before release, which was performed by 3 consecutive washes with warm PBS followed by incubation with DMEM + 10% FBS for 40-50 min. Cells were fixed and immunostained for tyrosinated α-tubulin. DNA was counterstained with DAPI. Only chromosomes spatially separated from the main mass of chromosomes were accounted as laggards.

### High throughput screening of lagging chromosomes

Quantification of lagging chromosomes was performed using 50-500 images of contiguous fields acquired in an IN Cell Analyzer 2000 microscope (GE Healthcare) with a Nikon 40x/0.95 NA Plan Fluor objective (binning 2×2), using a large chip CCD Camera (CoolSNAP K4) with a pixel array of 1024×1024 (2.7027 pixel/μm resolution). All early to late anaphase figures were classified regarding the presence or absence of lagging chromosomes. Any DAPI positive material between the two chromosome masses, but distinguishably separated from them, was accounted as lagging chromosomes. DNA bridges were excluded from this analysis.

### Photoactivation (PA) and photoconversion (PC)

For PA and PC assays, both U2OS cells stably expressing PA-GFP-α-tubulin/mCherry-α-tubulin or mEos-α-tubulin respectively, were cultured on glass coverslips as described. Mitotic cells were identified by DIC microscopy and imaging was carried out using a Plan-Apo 100× NA 1.40 DIC objective on a Nikon TE2000U inverted microscope equipped with a Yokogawa CSU-X1 spinning-disc confocal head containing two laser lines (488 nm and 561 nm) and a Mosaic (Andor) photoactivation system (405 nm). Photoactivation and photoconversion were carried out in late prometaphase or metaphase cells, the latter treated with 10 μM MG132 for less than 2 h, identified by mCherry-α-tubulin or mEos-α-tubulin signal, respectively. A region of interest for PA/PC was selected using a line segment placed perpendicular to the main axis in one of the sides adjacent to the metaphase plate. Seven 1 μm separated z-planes centered in the middle of the mitotic spindle were captured every 10-15 sec for 4.5 min (a pre-PA/PC image was always acquired) using 561 nm and 488 nm lasers and an iXon+ EM-CCD camera. Microtubule half-life was calculated using a MATLAB algorithm for kymograph generation and analysis. Briefly, generated kymographs were collapsed and the fluorescence intensity curve was created. Fluorescence intensity peaks of each time frame were defined by manually tracking regions of the highest fluorescence intensities followed by automated curve fitting and normalized intensities to the first time-point after PA/PC following background subtraction (background values obtained from quantifying the respective non-activated half-spindle). Values were corrected for photobleaching by normalizing to the values obtained from the quantification of fluorescence loss of whole-cell sum projected images. Microtubule turnover was calculated based on a fitted curve of the normalized intensities at each time point (corrected for photobleaching) to a double exponential curve A1*exp(−k1*t) + A2*exp(−k2*t) using MATLAB (Mathworks), in which *t* is time, A1 represents the less stable (non-kMTs) population and A2 the more stable (kMT) population with decay rates of k1 and k2, respectively (cells displaying an R squared value <0.99 were excluded from quantification). From the curve, the rate constants and the percentage of microtubules for the fast (typically interpreted as the fraction corresponding to non-kMTs) and the slow (typically interpreted as the fraction corresponding kMTs) processes. The half-life was calculated as ln2/k for each microtubule population.

### Tracking of centrosome separation

Detailed quantitative analysis of centrosome location and nuclear envelope topology was performed using custom made MATLAB scripts (The MathWorks Inc., USA; R2018a). Tracking of centrosome position/trajectories was performed in three-dimensional (3D) space using image stacks with a pixel size of 0.190 μm and z-step of 0.7 μm. Nuclear shape was reconstructed in 3D using H2B-GFP as a marker. From the centrosome locations and nuclear envelope reconstruction, it was possible to calculate the angle between the centrosome-centrosome axis and the nucleus major axis at NEB.

### Micropatterning

10 μm-width line micropatterns to control individual cell shape and adhesion pattern were produced as follows: glass coverslips (22 × 22mm No. 1.5, VWR) were activated with plasma (Zepto Plasma System, Diener Electronic) for 1 min and incubated with 0.1 mg/ml of PLL(20)-g[3,5]-PEG(2) (SuSoS) in 10 mM HEPES at pH 7.4, for 1 h, at room temperature. After rinsing and air-drying, the coverslips were placed on a synthetic quartz photomask (Delta Mask), previously activated with deep-UV light (PSD-UV, Novascan Technologies) for 5 min. 3 μl of MiliQ water were used to seal each coverslip to the mask. The coverslips were then irradiated through the photomask with the UV lamp for 5 min. Afterwards, coverslips were incubated with 25 μg/ml of fibronectin (Sigma-Aldrich) and 5 μg/ml of Alexa 647-conjugated fibrinogen (Thermo Fisher Scientific) in 100 mM NaHCO3 at pH 8.6, for 1 h at room temperature. Cells were seeded at a density of 50.000 cells/coverslip and allowed to spread for ~10-15h before imaging. Non-attached cells were removed by changing the medium ~2h-5h after seeding.

### Western-blotting analysis

Cells were grown until 90% confluence in complete growth medium and harvested by centrifugation at 1200 rpm for 5 min. Cell pellets were washed once with warm PBS and re-suspended in ice-cold lysis buffer (50 mM Tris HCl pH 7.4, 150 mM NaCl, 1 mM EDTA, 1 mM EGTA, 0.5% NP40, and 0.5% Triton™ X-100) supplemented with a cocktail of protease inhibitors (Roche). Protein samples were denatured in Laemmli buffer at 95°C for 5 min and 50 μg of total protein were separated by 10% (v/v) SDS-PAGE electrophoresis. Proteins were transferred to a nitrocellulose membrane using an IBlot™ Dry Blotting System (Invitrogen™). TTL and tubulins were probed using the following antibodies: rabbit polyclonal anti-TTL (1:2000, ProteinTech), rat monoclonal anti-tyrosinated α-tubulin clone YL1/2 (1:2000, Bio-Rad), rabbit polyclonal anti-detyrosinated α-tubulin(Liao et al., 2019), mouse monoclonal anti-polyglutamilated tubulin (1:1000, Adipogen), mouse monoclonal anti α-tubulin B-5-1-2 clone (1:10000; Sigma-Aldrich) and mouse monoclonal anti-β-tubulin clone T5201(1:2000, Sigma-Aldrich). Other proteins were immunodetected using mouse monoclonal anti-Cas9 (1:2500, Merck), anti-mCherry (1:2500, kind gift from I. Cheeseman, MIT, Cambridge, MA, USA), rabbit or goat polyclonal anti-GFP (1:2500, made by our in-house facility and 1:1000; Rockland Immunochemicals Inc., respectively), rabbit polyclonal anti-MCAK (1:5000, kind gift from D. Compton, Geisel School of Medicine, Dartmouth, USA), mouse monoclonal anti-Myc (1:10000; Cell Signalling Technology) and mouse monoclonal anti-GAPDH (1:40000; Proteintech). HRP-conjugated secondary antibodies (1:5000-10000, Jackson Immunoresearch) were visualized using ECL system (Bio-Rad). Protein levels were quantified by chemiluminescence using a ChemicDoc™ XRS+ system (BioRad).

### Statistical Analysis

Logistic regression was performed to evaluate the impact of multiple treatments and the occurrence of lagging chromosomes, adjusting the model for the different experiments. Significance was set at α=0.05 and these analyses were performed using SPSS, version 24.0. Unpaired two-tailed Student’s *t* test was used to determine the significance of differences between two groups (fluorescence intensity, spindle length, inter-kinetochore distance and microtubule half-life time). The Student’s t test was adjusted to the significance of the variance between experiments. Significance was set at α=0.05 and these analyses were performed using SPSS, version 24.0, and Prism, version 7.0a.

