## Supplemental Figures for "Microtubule tyrosination/detyrosination specifies a mitotic error code"

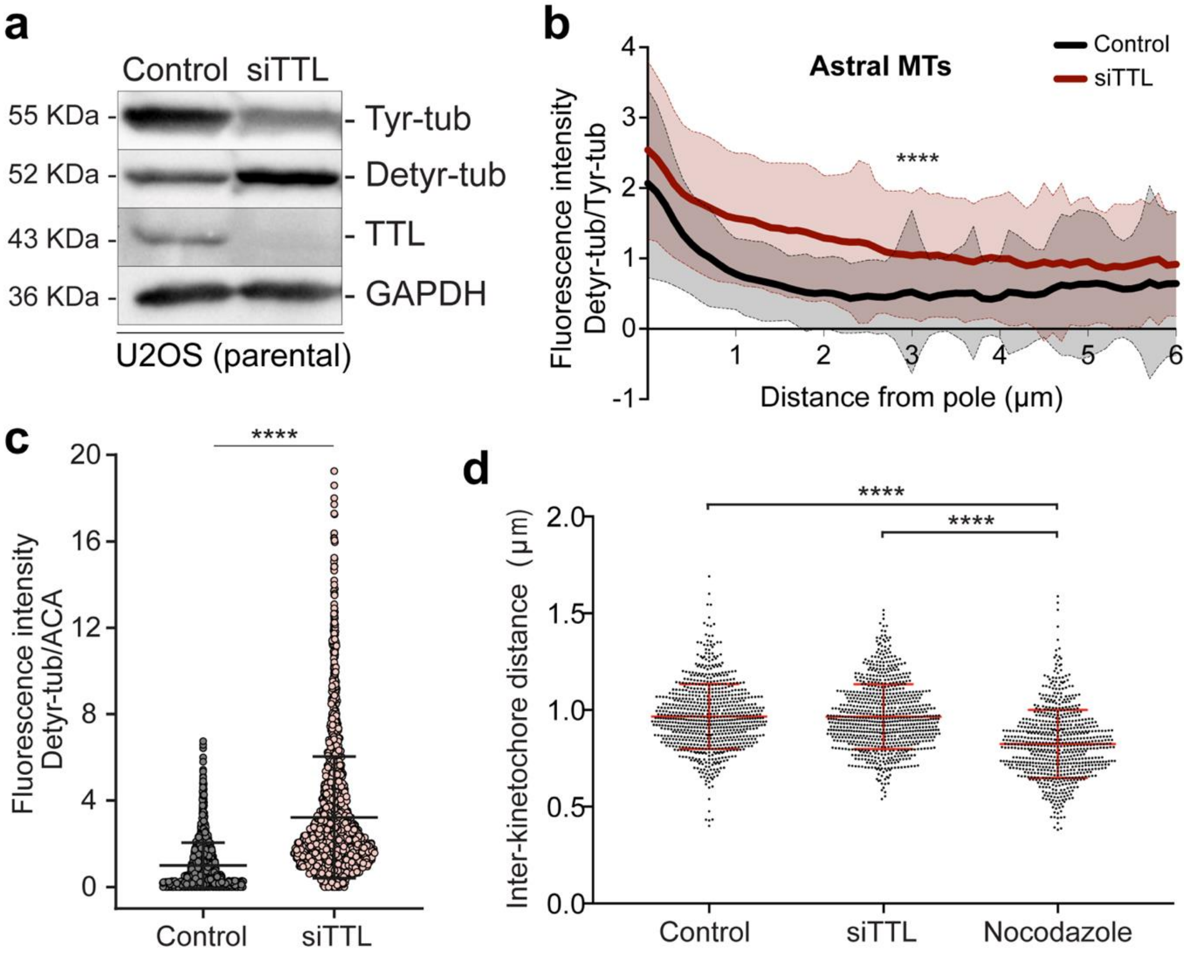


**Supplementary Figure 1│ TTL depletion increases α-tubulin detyrosination on astral and in the vicinity of kinetochores, without affecting inter-kinetochore distance.** (**a**) Protein lysates from parental U2OS cells collected 72h after RNAi transfection in control and after siTTL were immunoblotted for tyrosinated tubulin (Tyr-tub), detyrosinated tubulin (Detyr-tub) and TTL. GAPDH was used as loading control. (**b**) Quantification of the fluorescence intensity ratio between detyrosinated/tyrosinated α-tubulin along astral microtubules in control and siTTL metaphase cells (N=150 astral MTs/condition; 10 astral MTs/cell, 5 cells/experiment, pool of 3 independent experiments, $p<0.001$, unpaired two-tailed t-test). (**c**) Quantification of the fluorescence intensity of detyrosinated α-tubulin relative to ACA at the kinetochores (KTs) in control siTTL U2OS cells [N (Control) = 2224 KTs, 19 metaphase cells, pool of 2 independent experiments; N (siTTL)= 2507 KTs, 19 metaphase cells, pool of 2 independent experiments, $p<0.001$ (****), unpaired two-tailed t-test]. The mean fluorescence intensity of tubulin detyrosination at kinetochores normalized by the mean of control cells is represented. Error bars represent standard deviation. (**d**) Quantification of the inter-kinetochore distance in control, siTTL and nocodazole (1 μM) treated U2OS parental cells [N(control)=762 KT pairs, N(siTTL)=795 KT pairs and N(nocodazole)=695 KT pairs, from 6-11 cells/condition, pool of 3 independent experiments; $p<0.001$(****), unpaired two tailed t-test)].

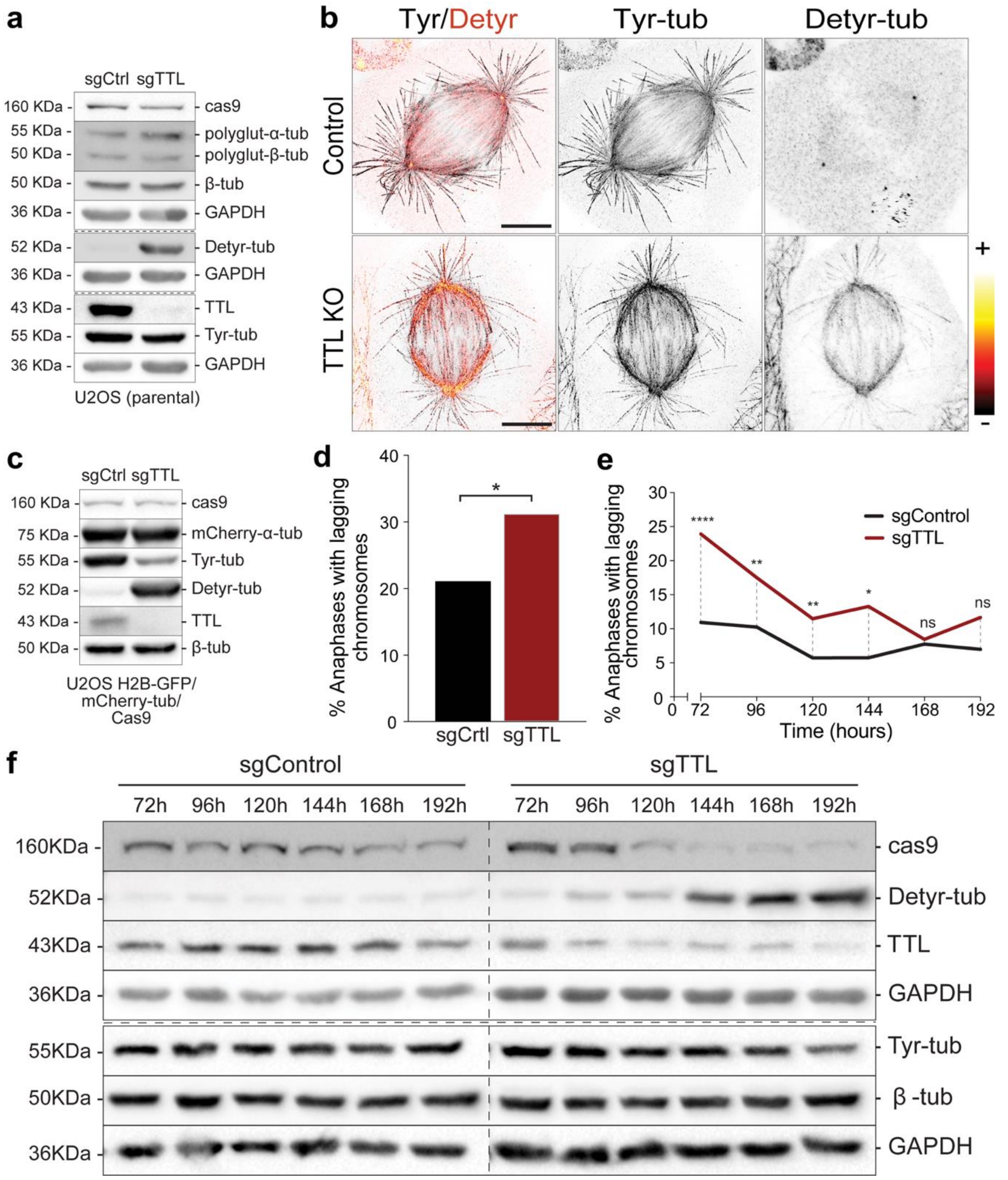


**Supplementary Figure 2│ Cellular adaptation to the chronic loss of TTL.** (**a**) Protein lysates of control (sgCtrl) and TTL KO (sgTTL) U2OS cells were immunoblotted for cas9, polyglutamylated tubulin (polyglut-α/β-tub), α-tubulin (α-tub), TTL, tyrosinated α-tubulin (Tyr-tub) and detyrosinated α-tubulin (Detyr-tub). GAPDH was used as loading control. (**b**) Confocal/CH-STED analysis of control (sgControl) and TTL KO (sgTTL) U2OS cells by immunofluorescence for Detyr-tub (confocal) and Tyr-tub (CH-STED). Red-hot lookup table was used to map extremes in fluorescence intensity for Detyr-tub. Scale bars, 5 µm. (**c**) Protein lysates of control (sgCtrl) and TTL KO (sgTTL) U2OS H2B-GFP/mCherry-tub cells were immunoblotted for cas9, mCherry, Tyr-tub, Detyr-tub and TTL. α-tubulin was used as loading control. (**d**) Quantification of the percentage of anaphase cells with lagging chromosomes in control (sgCtrl) and TTL KO (sgTTL) U2OS H2B-GFP/mCherry-tub cells by live-cell imaging between 72-168h after lentiviral transduction [N(sgCtrl) = 266 cells, N(sgTTL) = 241 cells, pool of 2 independent experiments, with 2 and 9 replicates, respectively, $p<0.05$(*), logistic regression]. (**e**) Quantification of the percentage of anaphase cells with lagging chromosome between 72-192h after lentiviral transduction in control (sgCtrl) and TTL KO (sgTTL) U2OS cells by immunofluorescence in fixed cells [N(sgCtrl)72h=324 cells, N(sgCtrl)96h=311 cells, N(sgCtrl)120h=280 cells, N(sgCtrl)144h=264 cells, N(sgCtrl)168h=320 cells, N(sgCtrl)192h=356 cells; N(sgTTL)72h=301 cells, N(sgTTL)96h=377 cells, N(sgTTL)120h=462 cells, N(sgTTL)144h=256 cells, N(sgTTL)168h=367 cells, N(sgTTL)192h=386 cells, pool of 3 independent experiments; $p<0.001$(****), $p<0.01$(**), $p<0.05$(*), ns=non-significant, unpaired two tailed t-test)]. (**f**) Protein lysates of control (sgControl) and TTL KO (sgTTL) U2OS cells were collected at different time points after lentiviral transduction and immunoblotted for cas9, Detyr-tub, TTL, Tyr-tub, and $\alpha$-tubulin. GAPDH was used as loading control.

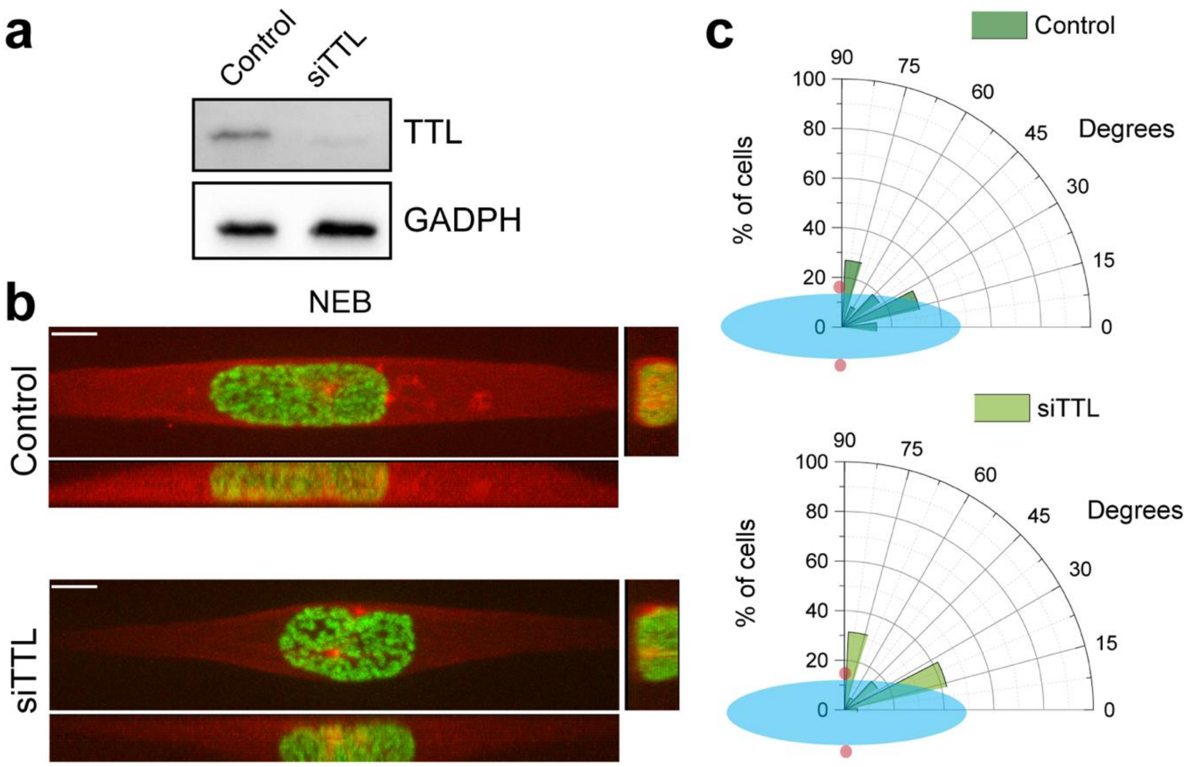


**Supplementary Figure 3│ The timing of centrosome separation at nuclear envelope breakdown is indistinguishable between control and TTL-depleted cells.** (**a**) Representative immunoblot to confirm TTL depletion efficiency by RNAi. (**b**) Live U2OS cells stably expressing H2B-GFP/mRFP-α-tubulin seeded on a horizontal, 10 μm-width line micropattern, at the moment of Nuclear Envelope Breakdown (NEB), in control and after siTTL. Scale bar, 10 μm. (**c**) Polar plot showing centrosome positioning relative to the long nuclear axis at NEB for control and siTTL cells [N(control)=22 cells, N(siTTL)=19 cells, pool of 3 independent experiments, non-significant differences between conditions, non-parametric Kolmogorov-Smirnov test). Nuclear shape is represented by a blue ellipse and centrosomes are represented by red circles.
